# The Gram-positive model organism *Bacillus subtilis* does not form microscopically detectable cardiolipin-specific lipid domains

**DOI:** 10.1101/190306

**Authors:** Alex-Rose Pogmore, Kenneth H. Seistrup, Henrik Strahl

## Abstract

Rather than being a homogenous diffusion-dominated structure, biological membranes can exhibit areas with distinct composition and characteristics commonly termed as lipid domains. Arguably the most comprehensively studied examples in bacteria are domains formed by cardiolipin, which have been functionally linked to protein targeting, cell division process, and mode of action of membrane targeting antimicrobials. Cardiolipin domains were originally identified in the Gram-negative model organism *Escherichia coli* based on preferential staining by the fluorescent membrane dye nonyl acridine orange (NAO), and later reported to exist also in other Gram-negative and -positive bacteria. Recently, the lipid-specificity of NAO has been questioned based on studies conducted in *E. coli*. This prompted us to re-analyse cardiolipin domains also in the Gram-positive model organism *Bacillus subtilis*. Here we show that logarithmically growing *B. subtilis* does not form microscopically detectable cardiolipin-specific lipid domains, and that NAO is not a specific stain for cardiolipin in this organism.

**Abbreviations:**

NAO: 10-nonyl acridine orange bromide
Nile Red: 9-diethylamino-5-benzo[±]phenoxazinone

## INTRODUCTION

Our understanding of the structure and organisation of biological membranes is based on the classical fluid mosaic model of Singer and Nicolson, which describes biological membranes as homogeneous two-dimensional fluids dominated by free lateral diffusion of lipids and embedded proteins [1]. However, the last decades of research have revealed that biological membranes are far more complex and heterogeneous than originally assumed, and specific lipid species can form distinct domains within biological membranes that fulfil specific biological functions [2, 3]. Arguably the most comprehensively studied type of bacterial lipid domain are clusters of cardiolipin forming at bacterial cell poles and cell division sites [4]. Cardiolipin is a complex phospholipid species commonly found in bacterial membranes, which is formed by two phosphatidylglycerol lipids linked together by an additional glycerol via phosphodiester bonds. Consequently, cardiolipin carries a double negative charge and a total of four fatty acid chains. What makes cardiolipin unique in the context of lipid domain studies is the fluorescent membrane dye nonyl acridine orange (NAO), which can be used to study the localisation and clustering-behaviour of cardiolipin in living bacterial cells [4, 5]. Due to its positive charge, NAO preferentially stains membranes containing anionic phospholipids. Upon interaction with cardiolipin, NAO undergoes a redshift in the fluorescent emission spectrum, thereby allowing the microscopic identification and visualisation of membrane areas that are enriched in cardiolipin [4, 5].

By using NAO-staining, Mileykovskaya and co-workers showed in their seminal study that *E. coli* membranes contain cardiolipin-enriched lipid domains that are localised at the cell poles and cell division sites [5]. Importantly, these findings were independently confirmed by analysing the composition of so-called minicells. These small cells are formed by a misplaced cell division occurring at the cell pole, which results in small anucleate cells that are highly enriched in cell material normally found at the cell poles. The analysis of the lipid composition of minicells revealed a clear enrichment of cardiolipin, thereby providing strong support for the polar cardiolipin domain hypothesis [6]. The mechanism through which cardiolipin accumulates at the cell poles is postulated to be based on a conical shape of the cardiolipin molecule, which provides preference for curved membrane found at the cell poles [7-9]. However, it should be noted that the shape of cardiolipin also depends on factors such as the type of fatty acids attached, pH and concentration of divalent cations, and that the ability to adopt a conical molecular shape is by no means unique to cardiolipin [10]. By forming a polar landmark that allows cardiolipin-interacting proteins to be specifically targeted to bacterial cell poles, cardiolipin domains have been suggested to be functionally linked to several prominent cellular processes such as cell division, chemotaxis, and transport [11-15]. Due to the high negative surface charge, polar cardiolipin domains have also been discussed as preferred targets for cationic membrane targeting antimicrobials such as host immunity peptides [16-19].

Recently, however, the specificity of NAO towards cardiolipin has been questioned [20]. By carefully analysing the staining-specificity both *in vitro* and *in vivo*, Oliver and co-workers concluded that the characteristic red-shifted fluorescence emission of NAO is not specific for dye molecules interacting with cardiolipin but rather promiscuous for anionic phospholipids in general. These studies were conducted in the Gram-negative model organism *E. coli* and much less attention has been given to other bacteria with respect to rigorous testing the specificity of NAO-staining. Instead, the broad conservation of cardiolipin-specific domains among bacterial species has been largely accepted within the community. Due to the concerns emerging from the *E. coli* studies, we decided to reanalyse both the existence of cardiolipin-domains, and the staining-specificity of NAO in the Gram-positive model organism *B. subtilis*. Here we show the domains previously observed in NAO-stained *B. subtilis* cells are a consequence of the used staining procedure, and are limited to certain media only. Furthermore, our data directly questions the specificity of NAO towards cardiolipin in *B. subtilis*, which has a significantly higher overall anionic phospholipid content than *E. coli* [21].

## METHODS

### Growth conditions

*Bacillus subtilis* cells were grown in either Lysogeny broth (LB) containing 5 g/l yeast extract (Oxoid), 10 g/l tryptone (Oxoid) and 10 g/l NaCl, or Difco sporulation medium (DSM) containing 8 g/l nutrient broth (Difco), 1 g/l KCl, 0.25 g/l MgSO_4_•7H_2_O, 1 mM Ca(NO_3_)_2_ 10 μM MnCl_2_, and 1 μM FeSO_4_. If required, the media was supplemented with tetracycline (5 μg/ml), kanamycin (2-5 μg/ml), spectinomycin (60 μg/ml) or erythromycin (2 μg/ml). The use of antibiotics was limited to strain construction only and all the experiments were carried out in the absence of selection pressure. In case of the stain KS60, both media were supplemented with 20 mM MgSO_4_, which is required for the growth of this strain.

For the microscopic experiments and the analysis of lipid content, the cells were grown overnight in the corresponding medium supplemented with 0.4% (w/v) glucose upon shaking at 30°C. The addition of glucose in the overnight cultures inhibits sporulation, thereby forming a more homogeneous starter culture. The overnight cultures were diluted 1:100 in the corresponding medium without glucose, and incubated upon shaking at 37°C until a logarithmic growth was obtained. The microscopy experiments were carried out with cells below OD_600_ of 0.5, the only exception being the cultures used for the analysis of lipid composition. In order to obtain sufficient cell material for this analysis, the cells grown in LB medium to an OD_600_ of 1.0.

### Strain Construction

As a widely used model organism, *B. subtilis* “wild type” strains have a long history of divergent laboratory evolution resulting in genetic and phenotypic variations. Instead of testing different strains from different laboratories for their ability to form cardiolipin domains, we chose to conduct our analysis with a wild type strain that is both well described and easy to obtain. For these reasons, all experiments shown in this manuscript are performed with *B. subtilis* 168 wild type obtained from Pasteur Institute, which is also the strain used to sequence and assemble the *B. subtilis* reference genome [22]. To exclude effects originating from the slightly different genetic backgrounds, all the deletions used in this study were transferred into the same wild type background though transformation with chromosomal DNA extracted from the respective donor strains [23]. See table 1 for strains and sources of chromosomal DNA used in this study.

**Table 1:** Strains used in this study.

| Strain | Relevant Genotype | Source | Construction |
| --- | --- | --- | --- |
| *B. subtilis* 168 | *trpC2* (wild type) | [22] | - |
| *B. subtilis* SDB206 | *clsA(ywnE)::ery*<br>*clsB(ywjE)::spc ywiE::neo* | [21] | - |
| *B. subtilis* HB5347 | *clsA(ywnE)::tet* | [37] | - |
| *B. subtilis* ARK3 | *clsA(ywnE)::tet clsB(ywjE)::spc*<br>*ywiE::neo* | this study | transformation of *clsA::tet*, *clsB::spc* and *ywiE::neo* into *B. subtilis* 168 |
| *B. subtilis* SDB01 | *psd::neo* | [36] | - |
| *B. subtilis* KS119 | *psd::neo* | this study | transformation of *psd::neo* into *B. subtilis* 168 |
| *B. subtilis* 4277 | Ωneo3427 Δ*mreB* *mbI::cat*<br>*mreBH::erm rsgI::spc* | [31] | - |
| *B. subtilis* KS60 | Ωneo3427 Δ*mreB* *mbI::cat*<br>*mreBH::erm rsgI::spc* | this study | transformation of Ωneo3427 Δ*mreB*, *mbI::cat*, *mreBH::erm*, and *rsgI::spc* into *B. subtilis* 168 |

### NAO and Nile Red membrane staining

The staining of *B. subtilis* with NAO was carried out with 200 μl culture aliquots in a round bottom 2 ml Eppendorf tubes with a perforated lid, and under constant shaking at 800 rpm with Eppendorf ThermoMixer. We have found this strict staining regime to reliably maintain adequate cell energisation in small culture volumes, thereby preventing the formation lipid domains linked to depolarisation-triggered delocalisation of MreB [24]. The time and temperature of the incubation varied depending on the specific experiment, and details are given in the context of the presented results. NAO (10-nonylacridine orange bromide, Sigma-Aldrich or Molecular Probes/Thermo Fisher Scientific) was solved in dimethyl sulfoxide (DMSO) and used at concentrations ranging from 100 nM to 2 μM. Upon staining, the concentration of solvent (DMSO) was maintained at 1% (v/v), which ensures good staining with hydrophobic membrane dyes without inhibiting cell growth or membrane integrity [25]. Nile Red (9-diethylamino-5-benzo[α]phenoxazinone, Sigma-Aldrich) solved in DMSO was added to culture aliquots at a final concentration of 500 nM (1% DMSO), followed by staining as described above for 5 min.

### Fluorescence Microscopy

Fluorescence microscopy was carried out with logarithmic growth phase cells grown under shaking at 37°C. The cells (0.5 μl culture aliquot) were immobilised on microscopy slides covered with a thin layer of 1.2% (w/v) agarose in H_2_O as described in detail elsewhere [25]. Fluorescence micrographs were captured with Nikon Eclipse Ti equipped with Photometrics Prime sCMOS camera, Nikon Plan Apo 100x/1.40 Oil Ph3 objective, and Sutter Instrument Company Lambda LS xenon arc light source. The excitation filters used is this study were ET470/40x and ET560/40x, the dichroic mirrors T495lpxr and T585lpxr, the emission filters ET525/50m and ET630/75m (Chroma). The NAO fluorescence in the green wavelength range was captured using the combination of ET470/40x, T495lpxr and ET525/50m. For the red NAO fluorescence, a combination of ET470/40x, T495lpxr and ET630/75m was used. The images were captured using Metamorph 7.7 (Molecular Devices) and analysed using Fiji [26].

### Fluorescence intensity measurements

The measurement of NAO fluorescence intensities at the green and red wavelength ranges was carried out with images exposed for 600 ms (525 nm) and 3000 ms (630 nm) in this order. The fluorescence intensities were measured from background-subtracted image fields along a 5 pixel wide and 50 pixel long line. For the quantification shown in Fig. 2 and Fig. S2, the analysed cell were chosen randomly based on phase contract images. For the quantitation shown in Fig. 1d-e, a cell depicted in Fig. 1a was chosen based on the highest apparent septal fluorescence signal. The intensity analysis was carried out with Fiji [26].

**Figure 1.**
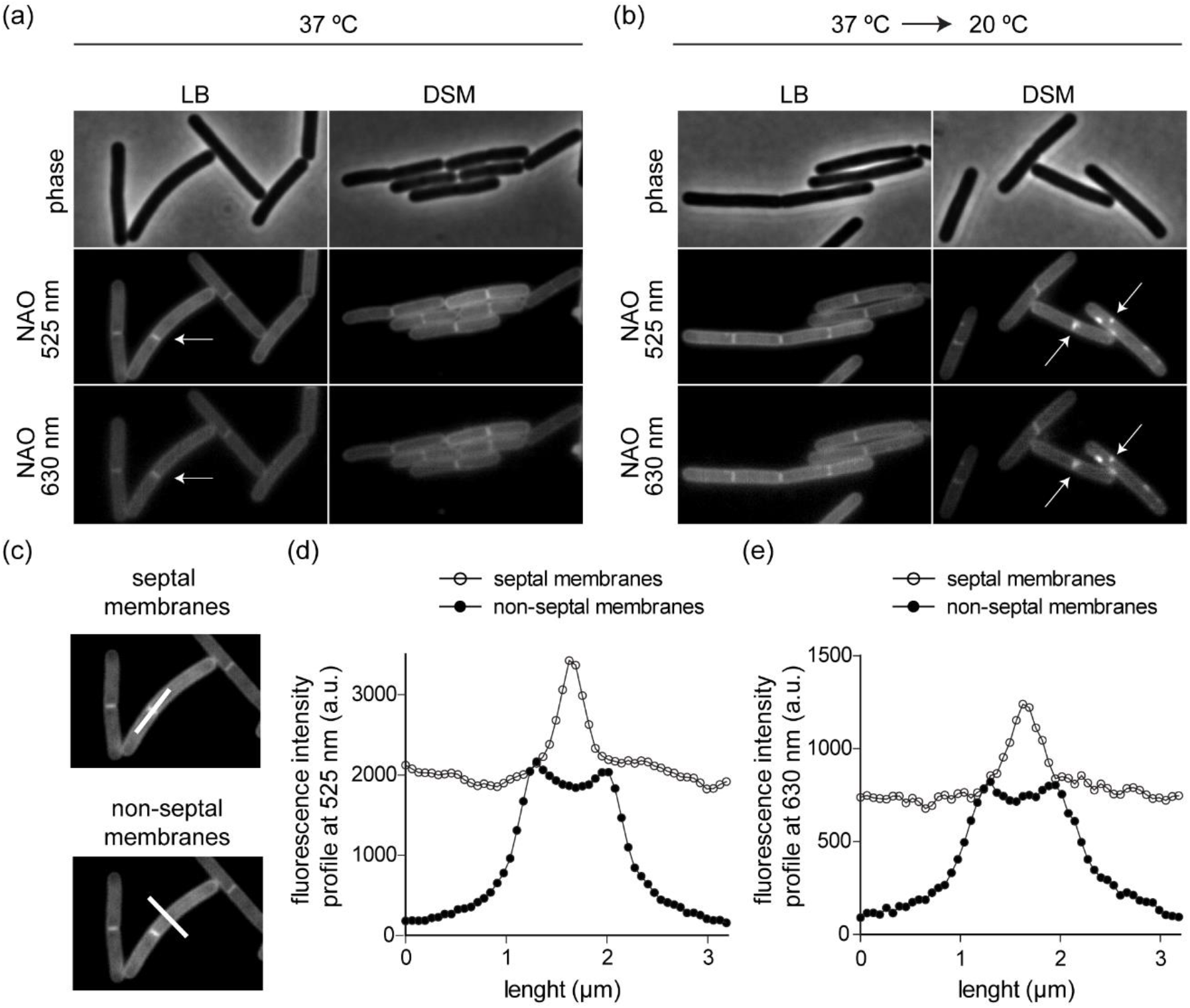
(a) Phase-contrast images (upper panels), and NAO fluorescent membrane stains at 525 nm (middle panels) and at 630 nm (lower panels) of logarithmic growth phase *B. subtilis* cells grown in LB and DSM-media at 37°C, respectively. For these images the cells were stained for 20 min at 37 °C. A cell division site (septum) is indicated with an arrow. The final concentration of NAO upon staining was 100 nM. (b) Comparable phase-contrast and fluorescent micrographs of *B. subtilis* cells grown at 37 °C, but stained with NAO for 20 min at 20°C. Few of the emerging domains are indicated with arrows. The final concentration of NAO upon staining was 100 nM. See supplementary Fig. 1 for a larger field of view with more cells. (c) Fluorescence micrographs of NAO-stained cells indicating the location of lines used for the fluorescence intensity measurement shown in panels d and e. (d-e) NAO fluorescence intensity line scans measured across the septum and across the length axis of the cell. The graphs depict the intensities at 525 nm (d) and the 630 nm (e), respectively. For this analysis, the cells were stained for 20 min at 37 °C. See supplementary Fig. S4 for the average intensity profiles of 10 analysed cells. Strain used: *B. subtilis* 168 (wild type)

**Figure 2.**
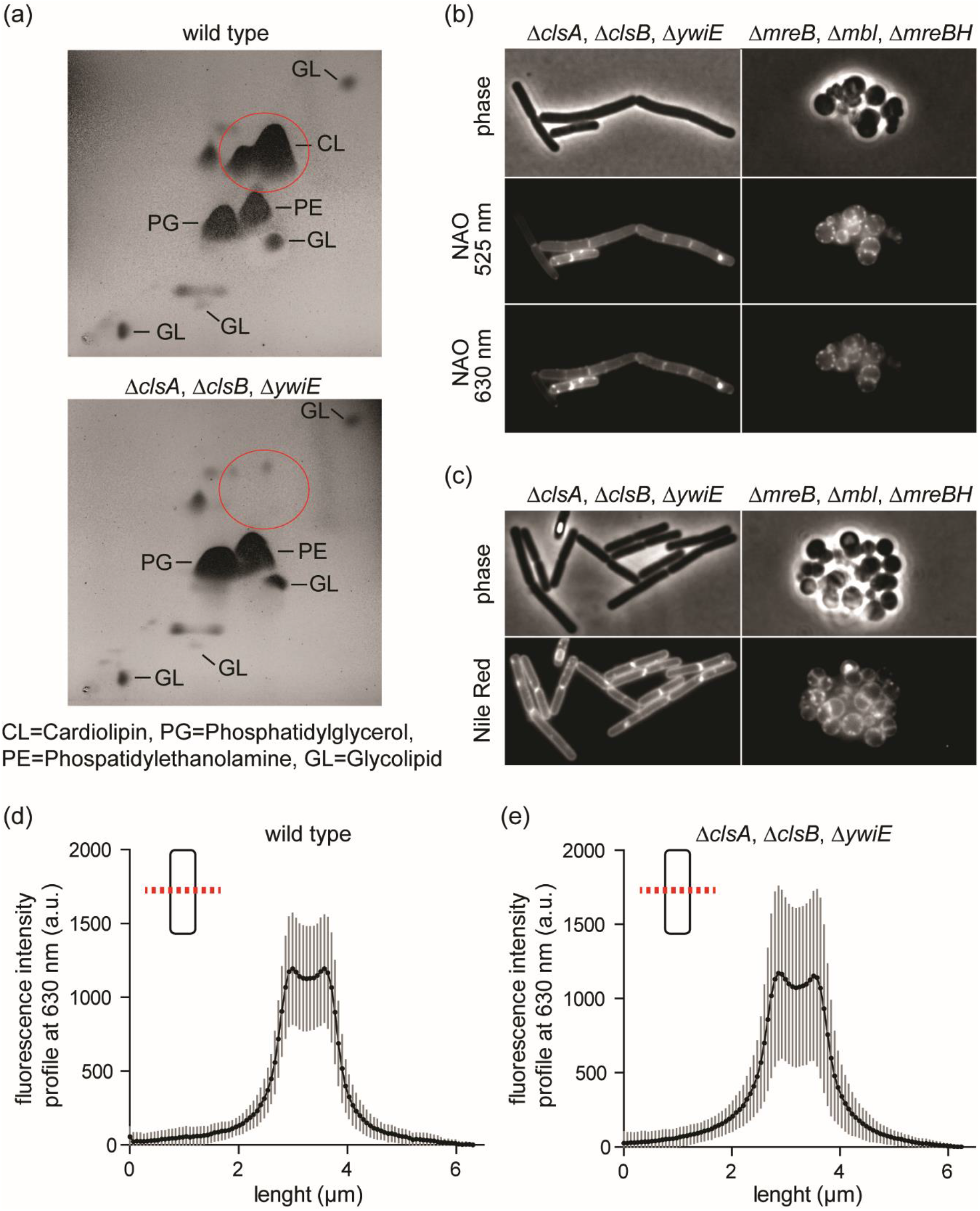
(a) Thin layer chromatography showing the phospholipid profiles of *B. subtilis* wild type, and *B. subtilis* strain deficient for cardiolipin synthase homologs. For the TLC-analysis, the cells were grown in LB medium at 37 °C, and harvested at OD600=1.0. (b) Phase-contrast images (upper panels), and NAO fluorescent membrane stains at 525 nm (middle panels) and at 630 nm (lower panels) of logarithmic growth phase *B. subtilis* cells deficient for cardiolipin synthase homologs (left panels) and >MreB homologs (right panels). The final concentration of NAO upon staining was 100 nM. (c) Phase-contrast images (upper panels), and Nile Red fluorescent membrane stains (lower panels) of logarithmic growth phase *B. subtilis* cells deficient for cardiolipin synthase homologs (left panels) and MreB homologs (right panels). The final concentration of Nile Red upon staining was 500 nM. The cells shown in panels (b) and (c) were grown in DSM medium at 37 °C, followed by staining for 20 min at 20 °C. In case of Nile Red, the dye was added only for the last 5 min of the 20 °C incubation period. See Fig. S5 and S6 for comparable micrographs of cells stained at the growth temperature (37 °C). (d/e) NAO fluorescence intensity profiles measured perpendicular to the cell length axis for *B. subtilis* wildtype cells (d), and for cells deficient for cardiolipin synthase homologs (e). Both graphs depict the average and standard deviations of profiles from 30 individual cells. Strains used: *B. subtilis* 168 (wild type), *B. subtilis* ARK3 (deficient for cardiolipin), and *B. subtilis* KS60 (deficient for MreB-homologs).

### Lipid Analysis

Phospholipid composition of *B. subtilis* 168 and ARK3 (Δ*clsA* Δ*clsB* Δ*ywiE*) was analysed by the Identification Service of the DSMZ, Braunschweig, Germany. For this aim, the strains were grown in LB at 37°C to an OD_600_ of 1, followed by harvest and wash in ice-cold 100 mM NaCl, and lyophilisation. The analysis at DSMZ was carried out by two dimensional silica gel thin layer chromatography (TLC) of lipids extracted using the Bligh and Dyer extraction method [27]. The TLC plates were stained using molybdatophosphoric acid, and the individual lipid species were identified using specific staining reagents as detailed in Tindall *et al* [28].

## RESULTS AND DISCUSSION

### Media and staining conditions promoting lipid domain formation

To our surprise, when we stained *B. subtilis* wild type cells grown in LB medium with low concentrations of NAO (100 nM), we did not observe the previously reported lipid domains (Fig. 1a). Instead, the NAO membrane stain was perfectly smooth both in the green wavelength range (500-550 nm), and in the potentially cardiolipin-specific red wavelength range (593-667 nm). The same absence of detectable lipid domains was observed in DSM medium, which was the medium used in the original publication describing cardiolipin domains in *B. subtilis* (Fig. 1a) [21]. In our group, the standard technique to stain cells with membrane dyes is to withdraw a 100-200 μl aliquot of an actively growing culture, and to stain the cell suspension in a round bottom 2 ml Eppendorf tube with a perforated lid at the growth temperature, and upon constant shaking. This is to ensure continuous aeration of the culture, which is crucial to maintain the cells adequately energised [25, 29], and to minimise changes in temperature, which can directly influence the analysed membranes by altering membrane fluidity and by triggering adaptation of the lipid composition [30, 31]. While the exact staining conditions are not comprehensively described in the original publication, the authors do mention that the staining was carried out for 20 min at room temperature [21]. To test if different staining temperatures could explain the discrepancy, we repeated the experiments with staining taking place at 20°C, as opposed to incubation at the growth temperature of 37°C. Indeed, under these conditions we could reproduce the previously published staining pattern with distinct domains present in the cytoplasmic membranes (Fig. 1b and S1). However, a frequent domain formation was only observed in cells grown in the DSM medium, and with a delay of approximately 20 min after the transfer to 20°C (Fig. S2). A large majority of cells grown in LB medium were free of NAO-stained lipid domains even upon staining at 20°C (Fig. 1b and S1). To rule out the possibility that absence of NAO-stained domains is due to batch variation of the dye, we compared staining with NAO purchased from two different manufacturers. In both cases, the membrane stains of cells grown in LB and stained at the growth temperature were perfectly smooth, thus ruling out this possibility (Fig. S3).

In the absence of cold shock, the only detectable local enrichment in fluorescent NAO-signal was the enhanced fluorescence emitted by the septum; a phenomenon commonly originating from close proximity of two parallel aligned septal membranes causing an increased apparent signal of any disperse membrane-associated fluorophore. To verify that this is indeed also the case for NAO, we quantified and compared the fluorescence intensities of NAO at septum, and at the lateral non-septal membranes (Fig. 2c and Fig. S4). It turned out that in both the green wavelength range (500-550 nm), and in the potentially cardiolipin-specific red wavelength range, the septal fluorescence intensities are in fact less than expected based on the presence of two septal membranes. The observed septal signal of NAO, thus, cannot be interpreted as enrichment of cardiolipin in the septal membranes. Consequently, we must conclude that actively growing *B. subtilis* cells do not form lipid domains that can be stained by NAO. Instead, the detected domains are a consequence of the staining procedure, and are limited to certain growth media.

### Cardiolipin-specificity of NAO in *B. subtilis* membranes

Next, we analysed the cardiolipin-specificity of the staining pattern observed in DSM medium upon cold shock. For this aim, we used a strain that carries deletions of the three known cardiolipin synthase genes *clsA, clsB* and *ywiE*. To verify that our strain indeed does not synthesise cardiolipin, we compared the thin layer chromatography profiles of lipid extracts from both the wild type and the cardiolipin synthase-deficient strain. As shown in Fig. 2a, the membranes of the tested cardiolipin synthase-deficient strain do not contain detectable levels of cardiolipin. Unexpectedly, the domain formation observed upon cold shock with NAO turned out to be indistinguishable between wild type cells and cardiolipin-deficient cells, thus arguing that the observed domains are not specific clusters of cardiolipin (Fig. 2a and Fig. S1). The existence of NAO-stained polar lipid domains in the absence of cardiolipin have also been reported for *E. coli* [20]. In this case, the domain formation was shown to require the synthesis of another common anionic lipid species phosphatidylglycerol. Hence, clustering of other negatively charged phospholipids that accumulate at the cell poles in the absence of cardiolipin was put forward as an explanation [20]. In contrast to *E. coli*, phosphatidylglycerol is an essential phospholipid species in *B. subtilis*, and depletion of phosphatidylglycerol synthase PgsA results in a lethal loss of membrane integrity [24]. Therefore, we chose to test the anionic nature of the observed lipid domains with an alternative method, and repeated the membrane staining of the cardiolipin-deficient deletion mutant with an uncharged fluorescent membrane dye Nile Red [32, 33]. The cold shock-triggered lipid domain pattern was readily detectable also with Nile Red (Fig. 2b and Fig. S5), thus ruling out a charge-driven mechanism for the preferential staining. While this experiment does not formally disprove the possibility that the observed domains are enriched in anionic phospholipids, negative charge does not appear to be their defining feature. Consequently, the cold-shock triggered domain formation in *B. subtilis* is most likely a phenomenon that is unrelated to the formation of polar anionic lipid domains in *E. coli*.

We have previously shown that dissipation of membrane potential results in delocalisation of bacterial actin homologs MreB, Mbl, and MreBH; a process that is linked to formation of fluid lipid domains that can be visualised with Nile Red [24, 29]. To test if the cold-shock induced lipid domains also depend on MreB-homologs, we repeated the membrane staining experiments in a strain that is deleted for *mreB*, *mbl*, and *mreBH*. However, these cells, which have lost their rod-shape due to the absence of MreB-dependent lateral cell wall synthesis [34], were still forming NAO and Nile Red-stained lipid domains in a cold shock-dependent manner (Fig. 2b-c and S6).

The mechanism that triggers the formation of the lipid domains observed upon cold shock, and the composition and physicochemical characteristic that define these domains remains elusive. More comprehensive future studies are needed to properly characterise this novel type of cold shock-triggered bacterial lipid domain. Localised clustering of cardiolipin, however, can be ruled out. At last, throughout our experiments we did not observe a noticeable difference in the intensity of the fluorescence membrane staining between cells that synthesise cardiolipin and cells that do not. This was also true for the red wavelength range, which has been postulated to be specific for cardiolipin (Fig. 2d-e). Consequently, we must conclude that, at least at the used concentration, NAO is not a valid microscopic reporter for cardiolipin in *B. subtilis*.

### Is the used concentration of NAO a relevant factor?

As mentioned earlier, Oliver and co-workers [20] provided conclusive evidence that the red-shifted fluorescence of NAO is not very specific for cardiolipin *in vitro*, and that polar NAO-stained lipid domains can be observed in *E. coli* even in the absence of cardiolipin. In addition, the authors state that they could reliably observe polar lipid domains only with NAO concentrations that are higher than what is commonly used [20]. To test if the lack of microscopically detectable NAO-stained lipid domains in *B. subtilis* is also linked to the used dye concentration, we stained wild type cells with different concentrations ranging from 100 nM to 2 μM. At higher concentrations of NAO domains that predominately localise to cell poles do indeed emerge (Fig. 3a). Worryingly, and unlike in *E. coli*, the concentration needed to observe these domains correlates with the minimal inhibitory concentration of NAO in *B. subtilis* (1 μM). Thus, it is questionable whether these domain are physiologically relevant. Irrespective of the possibility that the domains are in fact caused by NAO, the domain are also present in cells that do not synthesise cardiolipin (Fig. 3b), thus ruling out a cardiolipin-specific nature. Equally, the domains still emerge in the absence of MreB-homologs, thereby ruling out fluid lipid domain clustering caused by delocalisation of MreB as the primary cause [24].

**Figure 3.**
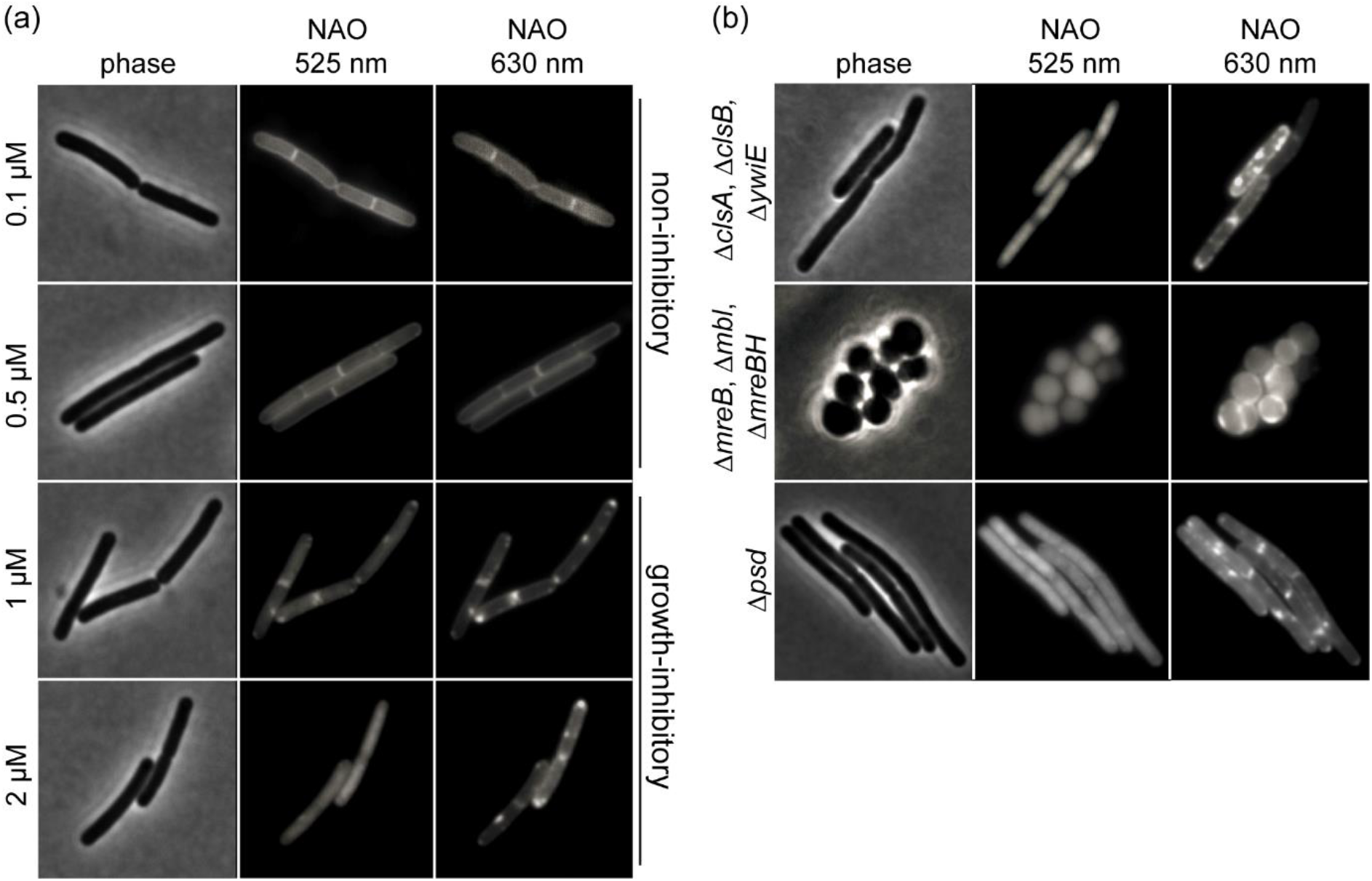
(a) Phase-contrast images (left panels), and NAO fluorescent membrane stains at 525 nm (middle panels) and at 630 nm (right panels) of logarithmic growth phase *B. subtilis* cells stained with varying concentrations of NAO (0.1-2 μM). Note that the increased concentrations of NAO (≥1 μM) results in detection of domains, an increasingly cytoplasmic signal in the green (525 nm) wavelength range, and in growth inhibition. (b) Phase-contrast images (left panels), and NAO fluorescent membrane stains at 525 nm (middle panels) and at 630 nm (right panels) of logarithmic growth phase *B. subtilis* cells stained with 2 μM NAO. Depicted is a comparison of strains deficient for cardiolipin (upper panels), MreB homologs (middle panels), and phosphatidylethanolamine (lower panels). The cells shown in this figure were grown in LB medium at 37 °C, followed by staining for 20 min at 37 °C. Strains used: *B. subtilis* 168 (wild type), *B. subtilis* ARK3 (deficient for cardiolipin), *B. subtilis* KS60 (deficient for MreB-homologs, and *B. subtilis* KS119 (deficient for phosphatidylethanolamine).

In principle, the domains observed with elevated concentrations of NAO in *B. subtilis* could be comparable to the anionic clusters identified by Oliver and co-workers in *E. coli* [20]. As mentioned earlier in the context of the cold-shock triggered domains, testing this possibility directly is difficult in *B. subtilis* since phosphatidylglycerol is an essential phospholipid in this organism. In *E. coli*, the main phospholipid species in the zwitterionic phosphatidylethanolamine, which can reach more than 70% of the total phospholipid content [35]. Consequently, anionic phospholipids can feasibly form distinct clusters in otherwise largely zwitterionic membrane environment. In *B. subtilis*, in contrast, phosphatidylethanolamine constitutes a smaller fraction, is not essential, and cells deficient for phosphatidylethanolamine accumulate anionic phosphatidylserine and glucolipids instead [36]. Crucially, these cells still exhibit the NAO-stained domains at elevated dye concentrations (Fig. 3b), indicating that the domains are unlikely to represent clusters of anionic phospholipids formed in a zwitterionic environment, as is the case in *E. coli*.

Comparable to the NAO-stained domains observed upon cold shock, the molecular identity of the domains observed with toxic concentrations of NAO, and perhaps even more importantly their physiological relevance remains an open question. What can be ruled out, however, is their identity as bona fide cardiolipin clusters.

### Summary

The experiments shown in this manuscript clearly demonstrate that NAO-fluorescence is not a reliable reporter for cardiolipin in the Gram-positive model organism *B. subtilis*. In the wider context, these findings highlight that, rather than relying on data obtained from other species, the potential lipid-specificity of NAO must be verified for the used bacterial species on a case-by-case basis, by using appropriate lipid synthase deletion strains. We would like to emphasise though, that our experiments by no means dismiss the possibility that the polar curved membranes of *B. subtilis* are enriched in cardiolipin. Rather, we simply lack reliable methods to detect potential clustering and local enrichment of cardiolipin.

## ACKNOWLEDGEMENT

We would like to acknowledge Kathi Scheinpflug for early microscopy experiments that led to this project.

## FUNDING INFORMATION

Funding for this project was provided by Newcastle University

## CONFLICTS OF INTEREST

The authors declare that there are no conflicts of interest

